# Environmental DNA (eDNA) metabarcoding surveys extend the range of invasion for non-indigenous freshwater species in Eastern Europe

**DOI:** 10.1101/2021.05.16.444374

**Authors:** Gert-Jan Jeunen, Tatsiana Lipinskaya, Helen Gajduchenko, Viktoriya Golovenchik, Michail Moroz, Viktor Rizevsky, Vitaliy Semenchenko, Neil J. Gemmell

## Abstract

Active environmental DNA (eDNA) surveillance through species-specific amplification has shown increased sensitivity in the detection of non-indigenous species (NIS) compared to traditional approaches. When many NIS are of interest, however, active surveillance decreases in cost- and time-efficiency. Passive surveillance through eDNA metabarcoding takes advantage of the complex DNA signal in environmental samples and facilitates the simultaneous detection of multiple species. While passive eDNA surveillance has previously detected NIS, comparative studies are essential to determine the ability of eDNA metabarcoding to accurately describe the range of invasion for multiple NIS versus alternative approaches. Here, we surveyed twelve sites, covering nine rivers across Belarus for NIS with three different techniques, i.e., an ichthyological, hydrobiological, and eDNA survey, whereby DNA was extracted from 500 mL surface water samples and amplified with two 16S rRNA primer assays targeting the fish and macro-invertebrate biodiversity. Nine non-indigenous fish and ten non-indigenous sediment-living macro-invertebrates were detected by traditional surveys, while seven NIS eDNA signals were picked up, including four fish, one aquatic and two sediment-living macro-invertebrates. Passive eDNA surveillance extended the range of invasion further north for two invasive fish and identified a new NIS for Belarus, the freshwater jellyfish *Craspedacusta sowerbii*. False-negative detections for the eDNA survey could be attributed to (i) preferential amplification of aquatic over sediment-living macro-invertebrates from surface water samples and (ii) an incomplete reference database. The evidence provided in this study recommends the implementation of both molecular-based and traditional approaches to maximize the probability of early detection of non-native organisms.

## Introduction

One of the main threats to native freshwater organisms is the establishment of and competition from non-indigenous species (NIS). In recent decades, this threat has intensified and accelerated through anthropogenic pressures, including climate change (Walther et al. 2009; Seebens et al. 2017). The introduction of NIS have the potential to transform local ecosystems through habitat transformation, community structure alteration, and evolutionary process modification (Mooney and Cleland 2001; Gallardo et al. 2019; Linders et al. 2019), causing economic consequences, negative impacts on ecosystem services and human well-being (Gallardo et al. 2019). Early detection is, therefore, essential for NIS management (Simberloff et al. 2005; Trebitz et al. 2017).

Two factors have facilitated the invasion of Ponto-Caspian species into Belarussian rivers, (i) the secondary connection of isolated river basins (Bij de Vaate et al. 2002) and (ii) global climate change (Semenchenko and Rizevskiy 2017). Many rivers in Belarus find their origin across the national border in two historically isolated basins, the Baltic Sea (e.g., Zapadnaya Dvina (Daugava) River, Neman River, Muchavets River) and Black Sea (e.g., Berezina River, Dnieper River, Pina River, Pripyat River, Sozh River) basins. To aid shipping transport within Belarus and throughout Europe, these two river basins have been secondarily connected via man-made canals (Bij de Vaate et al. 2002). The novel interconnectivity allowed Ponto-Caspian species to migrate north-westward into Belarus (Karatayev et al. 2008). Global climate change is generating water temperatures that facilitate the reproductive success of NIS, enhancing their spread into Belarus from Kiev (Ukraine), Kaunas (Lithuania) (Semenchenko and Rizevskiy 2017), and transboundary lakes and rivers.

Documenting the introduction and spread of NIS within Belarus commenced in the early 2000’s (Semenchenko et al. 2009). The current freshwater NIS checklist of Belarus includes 24 species of benthic macro-invertebrates (Lipinskaya et al., 2018; Semenchenko et al., 2016; Semenchenko et al., 2009) and 14 species of fish (Semenchenko and Rizevskiy 2017), with the majority of detections occurring in the southern part of the country (Semenchenko et al. 2016; Semenchenko and Rizevskiy 2017; Lipinskaya et al. 2018). The implemented monitoring techniques included standard hydrobiological and ichthyological surveys, with taxonomic identification through morphological characteristics (Karatayev et al. 2007; Semenchenko et al. 2009; Mastitsky et al. 2010). While species identification is feasible and easily obtainable for certain taxonomic groups (e.g., vertebrates), taxonomic identification through morphological characteristics is challenging for the majority of phyla, further complicated by the presence of juvenile and damaged specimens during collection. DNA-based technologies have, therefore, been implemented in recent years and helped to identify new non-native amphipod (*Echinogammarus trichiatus* (Martynov, 1932) (Lipinskaya et al. 2018)) and fish species (*Proterorhinus semilunaris* (Heckel, 1837) (Golovenchik et al. 2020).

Environmental DNA (eDNA), defined as intra- and extracellular DNA obtained directly from environmental samples (e.g., soil, sediment, water) without an obvious source of biological material (Taberlet et al. 2012), has been used in the last decade for the detection of species (Ficetola et al. 2008; Goldberg et al. 2013) and the investigation of ecological communities (Thomsen et al. 2012; Brett et al. 2016), including the early detection of non-indigenous (Dougherty et al. 2016; Ardura and Planes 2017; Hinlo et al. 2017; Klymus et al. 2017) and elusive (Piaggio et al. 2014; Simpfendorfer et al. 2016) species. Initially, the early detection of NIS through aquatic eDNA focused on active surveillance using targeted species-specific assays to assess the presence of a single species, achieving a higher detection probability and sensitivity then traditional monitoring approaches (Ardura et al. 2015; Dougherty et al. 2016; Simpfendorfer et al. 2016). However, active surveillance decreases in cost- and time-efficiency when multiple NIS are of interest (Rojahn et al. 2021).

Therefore, a shift towards passive NIS surveillance has occurred more recently (Holman et al. 2019; van den Heuvel-Greve et al. 2021), with eDNA metabarcoding taking advantage of the complexity of the DNA signal contained within environmental samples and enabling the simultaneous detection of multiple species (Cristescu 2014). Although eDNA metabarcoding has outperformed traditional ichthyological survey techniques in multiple studies (Hänfling et al. 2016; Cilleros et al. 2019), active surveillance through targeted amplification has shown increased detection sensitivity for rare species compared to eDNA metabarcoding (Harper et al. 2018; Bylemans et al. 2019). While NIS have been detected by eDNA metabarcoding (Holman et al. 2019; van den Heuvel-Greve et al. 2021), comparisons of this approach to traditional survey techniques are needed to determine the capability of passive eDNA surveillance to accurately describe the invasion range of non-indigenous species.

In this study, the range of invasion for fish and macro-invertebrates in Belarussian rivers was determined by three survey techniques, i.e., an ichthyological, hydrobiological, and eDNA metabarcoding survey. Our eDNA survey targeted two regions of the 16S rRNA gene for fish and crustacean detection. The number of NIS detected and the range of invasion of each NIS was compared between survey methods to determine the capability of eDNA metabarcoding to describe the invasion range of aquatic and sediment-living freshwater non-indigenous species in temperate riverine systems.

## Materials and methods

### Sampling sites

Twelve sites were sampled on nine water bodies across Belarus in May-June 2018 with three different monitoring methods to compare the detection efficiency of NIS between traditional survey techniques and eDNA metabarcoding (TABLE 1; FIGURE 1). These sites represent a subset of areas regularly monitored by the standard hydrobiological and ichthyological surveys conducted in Belarus for NIS documentation (Semenchenko et al. 2013). Sampling sites are characterized by different bottom structures and other environmental parameters (TABLE 1). Hydro-physical parameters (pH, conductivity, water temperature) were recorded by pH, EC/TDS, and Temperature Meters HANNA® HI 98311. The water pH varied during sampling from 6.8 to 8.5, conductivity from 210 μS to 396 μS, and temperature varied from 19.5 °C to 23.4 °C.

**Table 1.** Sampling sites and their description

| # | Cod<br>e | Water<br>body | Near<br>location | Latitud<br>e, N<br>(DD) | Longitud<br>e, E<br>(DD) | Type of<br>bottom<br>sedimen<br>ts | T,<br>°C | p<br>H | Conductivi<br>ty, µS |
| --- | --- | --- | --- | --- | --- | --- | --- | --- | --- |
| 1 | NZ | Dnieper<br>River | Nizhnie<br>Zhary<br>village | 51.296<br>16 | 30.57220 | Silty<br>sand | 23.<br>4 | 8.<br>5 | 285 |
| 2 | PN | Pripyat<br>River | Narovlya<br>town | 51.787<br>58 | 29.54408 | Silty<br>sand | 21.<br>2 | 7.<br>2 | 365 |
| 3 | PP | Pina<br>River | Pinsk town | 52.110<br>73 | 26.10597 | Pebble<br>and<br>sand | 23.<br>3 | 7.<br>7 | 396 |
| 4 | DB<br>D | Dnieper-Bug<br>Canal | Duboi<br>village | 52.081<br>47 | 25.78353 | Silt | 21.<br>2 | 8.<br>2 | 364 |
| 5 | DB | Dnieper-Bug<br>Canal | Vigoda<br>village | 52.148<br>13 | 24.71443 | Silt | 21.<br>1 | 7.<br>3 | 210 |
| 6 | B | Muchave<br>ts River | Brest town | 52.087<br>65 | 23.72958 | Silt | 21.<br>5 | 7.<br>8 | 210 |
| 7 | S | Sozh<br>River | Chenki<br>village | 51.787<br>58 | 30.94556 | Silt | 22.<br>0 | 6.<br>8 | 327 |
| 8 | DM | Dnieper<br>River | Mogilev<br>town | 53.980<br>54 | 30.39546 | Silty<br>sand | 21.<br>1 | 8.<br>1 | 280 |
| 9 | BZ | Berezina<br>River | Panjushkovi<br>chi vill. | 53.235<br>02 | 29.18072 | Silty<br>sand | 21.<br>8 | 8.<br>5 | 330 |
| 10 | N | Neman<br>River | Gozha<br>village | 53.809<br>05 | 23.85000 | Sand | 22.<br>4 | 7.<br>8 | 200 |
| 11 | DV<br>D | Zapadna<br>ya Dvina<br>River | Verchnedvin<br>ks town | 55.744<br>71 | 27.92473 | Silt | 19.<br>5 | 8.<br>0 | 266 |
| 12 | ZD | Zapadna<br>ya Dvina<br>River | Druja town | 55.793<br>53 | 27.44047 | Silt | 19.<br>5 | 8.<br>0 | 268 |

**Figure 1:**
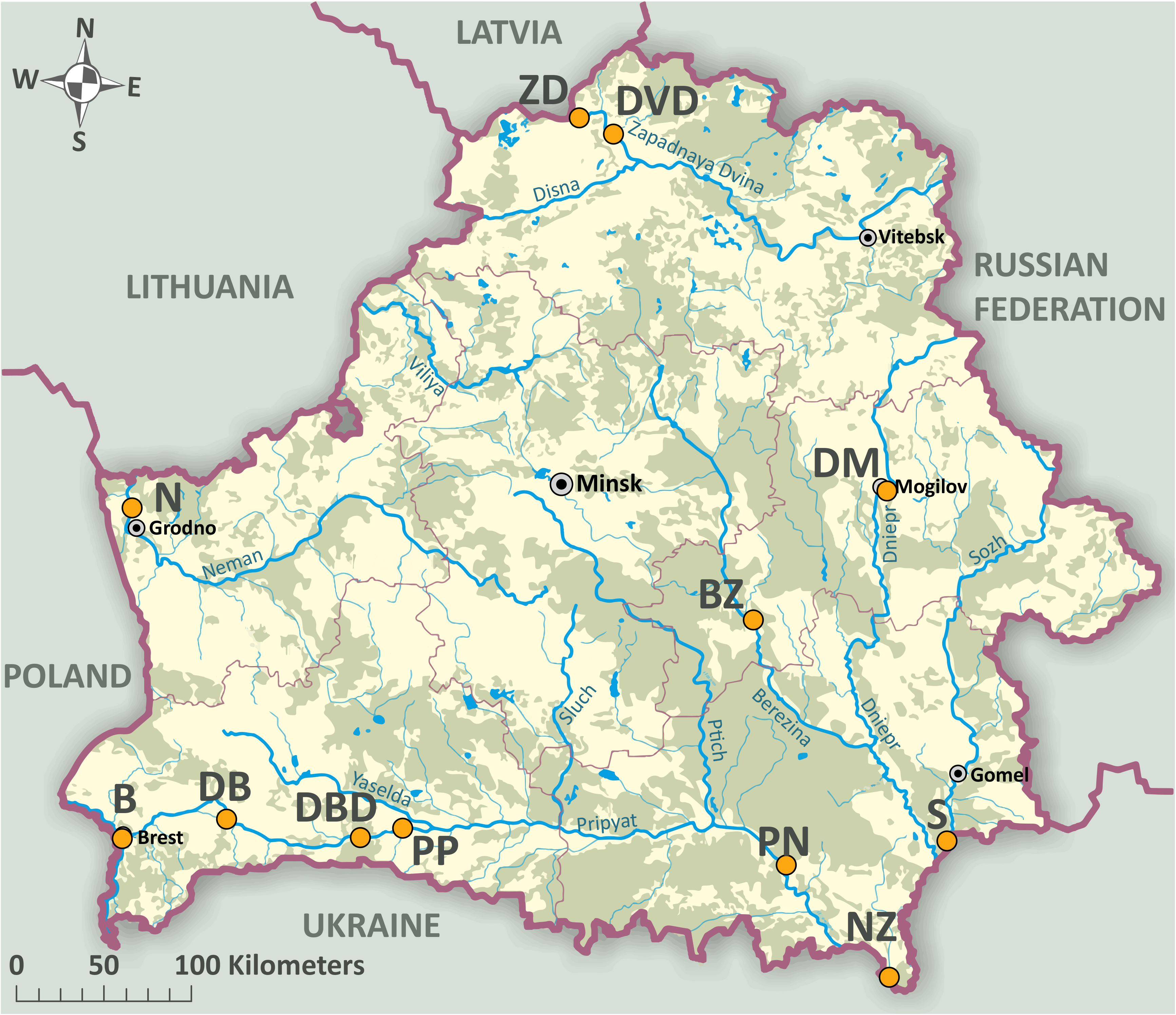
Map of Belarus displaying the twelve sampling sites. Sampling sites are indicated by orange-colored circles. Sampling site notation follows the abbreviations of Table 1.

### Hydrobiological survey

Quantitative and qualitative benthic macro-invertebrate samples were taken by hand-net (ISO 7828; 25 cm × 25 cm frame; 500 μm mesh size) at each of the twelve sites. Two macro-invertebrate samples were obtained from the littoral zone of each site at a depth of 50 – 70 cm. For quantitative assessment, samples were collected by pushing the hand-net gently through the uppermost 2 – 5 cm of the substratum and dragging it for 3-5 m. For qualitative assessment, multiple smaller samples were collected from different habitats of the sample site to maximize the diversity of captured taxa. Samples were fixed in 96% ethanol and sorted in the laboratory. Specimens were identified to the lowest possible taxonomic level using identification keys, resulting in higher taxonomic ranks for certain groups, i.e., Hydrachnidia, Oligochaeta, and Diptera. Moreover, juvenile and damaged specimens from the taxonomic groups of Mollusca, Ephemeroptera, and Coleoptera were identified to the genera or family level.

### Ichthyological survey

Two survey techniques were employed for the ichthyological survey dependent on the sampling site, seining (30 m length, 8 – 10 mm mesh size) and hand netting (60 cm × 55 cm frame; 8 mm mesh size). Ichthyological surveys were conducted in the littoral shallow part of the sampling sites. Species identification through morphological characteristics occurred on site. Native fish species were released back into their habitat upon identification, while non-indigenous species were collected for ichthyological and genetic purposes to the Laboratory of Ichthyology, Scientific and Practical Center for Bioresources, National Academy of Sciences of Belarus, Minsk, Belarus. Individual counts per species were used to infer abundance.

### Environmental DNA survey

Aquatic eDNA sampling was performed concurrent to the hydrobiological survey. Within each of the twelve sites, nine surface water samples were collected covering three habitats, with three biological replicate samples per habitat. Sampling occurred from 30 May until 10 June 2018. Environmental DNA filtration followed recommendations from (Spens et al. 2016). Briefly, pre-packed sterile 50 mL luer-lock syringes were used to push the sampled water through the Sterivex™ (Millipore, Merck KGaA, Darmstadt, Germany) column until clogging. The volume of water filtered through Sterivex™ columns ranged from 250 mL to 750 mL, depending on the turbidity of the water column. Mean volume of water per filter equaled 424.3 ± 124.8 mL (± SD), while mean volume of water per site equaled 3,641.67 ± 982.77 mL (± SD). Remaining water in the columns was removed by pushing air through the filter. Pre-packed sterile 5 mL luer-lock syringes were used to add 2 mL Longmire’s Buffer (100 mM Tris, pH 8.0; 100 mM EDTA, pH 8.0; 10 mM NaCl; 0.5% Sodium Dodecyl Sulfate; 0.2% Sodium Azide) to each column. Samples were stored at −20 °C until shipment on ice to the eDNA facility at the University of Otago, Dunedin, New Zealand, where samples were stored at −20 °C until further processing.

### DNA extraction

Prior to laboratory work, all bench surfaces and equipment were sterilized by a 10 minute exposure to 10% bleach solution (Prince and Andrus 1992) and rinsed with ultrapure water. To test for contamination, negative filtration controls (500 mL ultrapure water), negative extraction controls (500 μL ultrapure water), and negative PCR controls (2 μL ultrapure water) were added and processed alongside the samples.

Sample processing followed the recommendations in (Spens et al. 2016) with slight modifications. Briefly, DNA was extracted solely from the Longmire’s Buffer, since DNA extracts from filters did not show amplification success during initial testing. Caps were taken off Sterivex™ columns and buffer was transferred to a 2 mL Eppendorf LoBind tube using a pre-packed sterile 5 mL luer-lock syringe. Samples were spun at 6000 × g for 45 minutes, after which the supernatant was discarded. 180 μL ATL and 20 μL proteinase K were added to the pellet. Samples were briefly vortexed and incubated at 56 °C overnight in a spinning rotor. Following 15 seconds of vortexing, equal volumes of buffer AL and 100% ethanol were added to the sample. After mixing, the standard protocol of the Qiagen DNeasy Blood & Tissue Kit (Qiagen GmbH, Hilden, Germany) was followed. DNA extracts were stored at −20 °C until further processing.

### Library preparation

Library preparation followed the protocol described in (Jeunen et al. 2018). Briefly, two metabarcoding assays targeting two fragments of the 16S rRNA gene region were used to amplify DNA from fish and crustaceans (SUPPLEMENT 1). Prior to library preparation, the presence of inhibitors was tested for and low-template samples were identified by a dilution series (neat, 1/10, 1/20). Amplification was carried out in triplicate in 25 μL reactions and qPCR mastermix consisted out of 1x SensiFAST SYBR^®^ Lo-ROX Mix (Bioline, London, UK), 0.4 μmol/L of each primer (Integrated DNA Technologies, Australia), 2 μL of template DNA, and ultrapure water as required. The thermal cycling profile included an initial denaturation step at 95 °C for 10 minutes; then 50 cycles of 30 seconds at 95 °C, 30 seconds at 51-54 °C (see annealing temperatures in SUPPLEMENT 1), 45 seconds at 72 °C; and a final extension of 10 minutes at 72 °C.

A one-step amplification protocol using fusion primers was employed for library building. Fusion primers contained an Illumina adapter, a modified Illumina sequencing primer (absent in the reverse fusion primer), a barcode tag (6-8 bp in length), and the template specific primer. Each sample was amplified in duplicate and assigned a unique barcode combination. qPCR conditions followed the protocol described for the inhibition test. Post qPCR, sample duplicates were pooled to reduce stochastic effects from PCR amplification. Samples were then pooled based on end-point qPCR fluorescence and Ct-value in mini pools. Size selection and qPCR clean-up followed the AMPure XP (Beckman Coulter, US) standard protocol. Molarity of mini pools was measured on Agilent 2100 Bioanalyzer (Agilent, The Netherlands) and pooling occurred equimolarly to produce a single DNA library. Final concentration of the library was assessed via Qubit. Sequencing was performed by Otago Genomics on Illumina MiSeq^®^ (300 cycle, single-end V2 kit), following the manufacturer’s protocols, with 10% of PhiX to minimize issues associated with low-complexity libraries.

### Bioinformatic and statistical analyses

The bioinformatic analysis for both assays followed an in-house bioinformatic pipeline using FastQC v.0.11.5 (Bioinformatics 2011), Geneious Prime v.11.0.3+7 (Kearse et al. 2012), VSEARCH v.2.13.3 (Rognes et al. 2016), and OBITools v.1.2.11 (Boyer et al. 2016). Raw fastq files were checked for quality using FastQC. Reads were separated by barcode and assigned to samples using the ‘separate reads by barcode’ function in Geneious Prime, allowing for a single mismatch. All barcodes had a minimum three basepair mismatch distance from each other. Primer sequences were removed, allowing for a single mismatch, using the ‘annotate new trimmed regions’ function in Geneious Prime. The remaining reads were exported in fastq format and subsequently filtered based on total expected errors “-- fastq_maxee 0.1”, minimum length “--fastq_minlen 100”, maximum length “--fastq_maxlen 230”, and ambiguous bases “--fastq_maxns 0”, using the ‘--fastq_filter’ function in VSEARCH. Successful quality filtering was checked by FastQC report. The remaining sequences were dereplicated into unique sequences using the ‘--derep_fulllength’ function and unique sequences with an abundance lower than 50 were removed. Unique sequences were clustered at 97% using the ‘--cluster_size’ function, followed by the removal of chimeric sequences with the function ‘--uchime3_denovo’. Finally, an OTU table was generated at 97% threshold using the ‘--usearch_global’ function.

All OTUs were assigned a taxonomy using the ‘ecotag’ function in OBITools. A custom reference database was generated for both metabarcoding assays by an *in silico* PCR using the ‘ecoPCR’ function on the EMBL dataset (downloaded on the 13^th^ of May 2020). The custom reference database was supplemented with the newly barcoded NIS sequences. Further filtering was conducted on the taxonomic assignment table prior to statistical analysis. All OTUs failing to obtain a taxonomic assignment were discarded from the dataset, as well as unspecific taxonomic targets, a positive detection in a sample represented by a single sequence and OTUs with positive detections in negative control samples. Finally, identical taxonomic assignments were summed per sample and replicates per site were summed to obtain a single taxonomic list per sampling site.

To assess the proportion of missing reference sequences and issues surrounding amplification efficiency from mismatches in primer-binding sites, the custom reference database generated for both metabarcoding assays by the *in silico* PCR (‘ecoPCR’) function was checked for the presence of NIS detected by both traditional survey methods. Rarefaction curves were generated to assess sequencing coverage using the ‘rarecurve’ function from the ‘vegan v.2.4-1.’ package in R v.3.3.2 (R; http://www.R-project.org).

## Results

### Biodiversity detection

A total of 43 fish species were identified across the twelve sampling sites with our ichthyological survey, representing twelve families and eight orders, including Cypriniformes, Perciformes, Syngnathiformes, Osmeriformes, Gadiformes, Siluriformes, Clupeiformes, and Salmoniformes (SUPPLEMENT 2).

Our hydrobiological survey identified a total of 133 macro-invertebrate taxa across all twelve sampling sites, covering 66 families and four phyla, i.e., Cnidaria, Mollusca, Annelida, and Arthropoda (SUPPLEMENT 3). Taxonomic assignment through morphological characteristics allowed us to identify 97 taxa to species level, while 19 taxa were identified to genus and 17 taxa to family level.

Filtering and quality control returned 5,661,054 reads, with 3,845,772 and 1,815,282 reads for the fish (16S) and crustacean (16S) metabarcoding assays, respectively. Overall, eDNA samples achieved good sequencing coverage, based on rarefaction curves (SUPPLEMENT 4), with a mean number of reads per habitat ± s.d.: fish (16S): 113,662 ± 137,606; crustacean (16S): 51,932 ± 35,319. Amplification difficulties, resulting in low coverage, were encountered in sample 3-N for the fish (16S) assay. This sample was removed from the final dataset, prior to analysis. A total of 31 and 246 OTUs were recovered for the fish (16S) and crustacean (16S) metabarcoding assays, respectively. Further stringent quality control post taxonomic assignment reduced the number of total detections to 15 and 75, covering 8 and 36 families for the fish (16S) and crustacean (16S) metabarcoding assays, respectively (SUPPLEMENT 5).

### Non-indigenous species detection

Nine non-indigenous fish species were identified with our ichthyological survey, all of which have previously been detected in Belarus, including racer goby (*Neogobius gymnotrachelus*), monkey goby (*Neogobius fluviatilis*), western tubenose goby (*Proterorhinus semilunaris*), Chinese sleeper (*Perccottus glenii*), black-striped pipefish (*Syngnathus abaster*), southern nine spine stickleback (*Pungitius platygaster*), stellate tadpole-goby (*Benthophilus stellatus*), common carp (*Cyprinus carpio carpio*), and Back Sea sprat (*Clupionella cultriventris*).

Ten out of 24 established non-indigenous macro-invertebrates were detected by the 2018 hydrobiological survey. These include one mysid (*Limnomysis benedeni*), six amphipods (*Chelicorophium curvispinum*, *Chelicorophium robustum*, *Dikerogammarus haemobaphes*, *Echinogammarus ischnus*, *Obesogammarus crassus*, and *Obesogammarus obesus*), one decapod (*Faxonius limosus*), and two invasive alien mollusks (*Lithoglyphus naticoides* and *Dreissena polymorpha*). Both alien mollusks were excluded in the comparative analysis, as the phylum Mollusca is not amplifiable by the primer assays used in our eDNA survey.

Our eDNA metabarcoding survey detected seven non-indigenous species, including four fish and three macro-invertebrate species. All four non-indigenous fish species were detected by our ichthyological survey, i.e., racer goby (*Neogobius gymnotrachelus*), western tubenose goby (*Proterorhinus semilunaris*), chinese sleeper (*Perccottus glenii*), and monkey goby (*Neogobius fluviatilis*). Two of the three non-indigenous macro-invertebrates were detected by the hydrobiological survey, i.e., spinycheek crayfish (*Faxonius limosus*) and *Dikerogammarus haemobaphes*. Our eDNA survey detected one additional NIS, a freshwater jellyfish (*Craspedacusta sowerbii*), not detected by the hydrobiological survey.

### Reference database analysis

Overall, eleven of the seventeen NIS detected by both traditional monitoring methods, including five out of nine fish and six out of eight macro-invertebrates, have a reference barcode in molecular databases for the 16S target region of our eDNA metabarcoding assays (FIGURE 2). The *in silico* PCR used to construct the reference database requires the presence of primer-binding sites, which are frequently removed prior to sequence depositing in the EMBL database when the same assay is used for barcoding. Therefore, two of the five non-indigenous fish were not picked up by the *in silico* PCR analysis, even though a reference for the amplicon is publicly available on the EMBL database. For this project, one non-indigenous fish (*Neogobius fluviatilis*, BOLD accession number 689-fB) and two non-indigenous macro-invertebrates (*Chelicorophium curvispinum*, BOLD accession number TLAMP475S-17; *Obesogammarus obesus*, BOLD accession number TLAMP330S-17) were barcoded. At the time of analysis, *Benthophilus setllatus*, *Clupionella cultriventris*, and *Cyprinus carpio carpio* do not have a reference barcode for the target region of the fish (16S) assay and, hence, cannot be identified, at least to species-level, by our eDNA metabarcoding survey.

**Figure 2:**
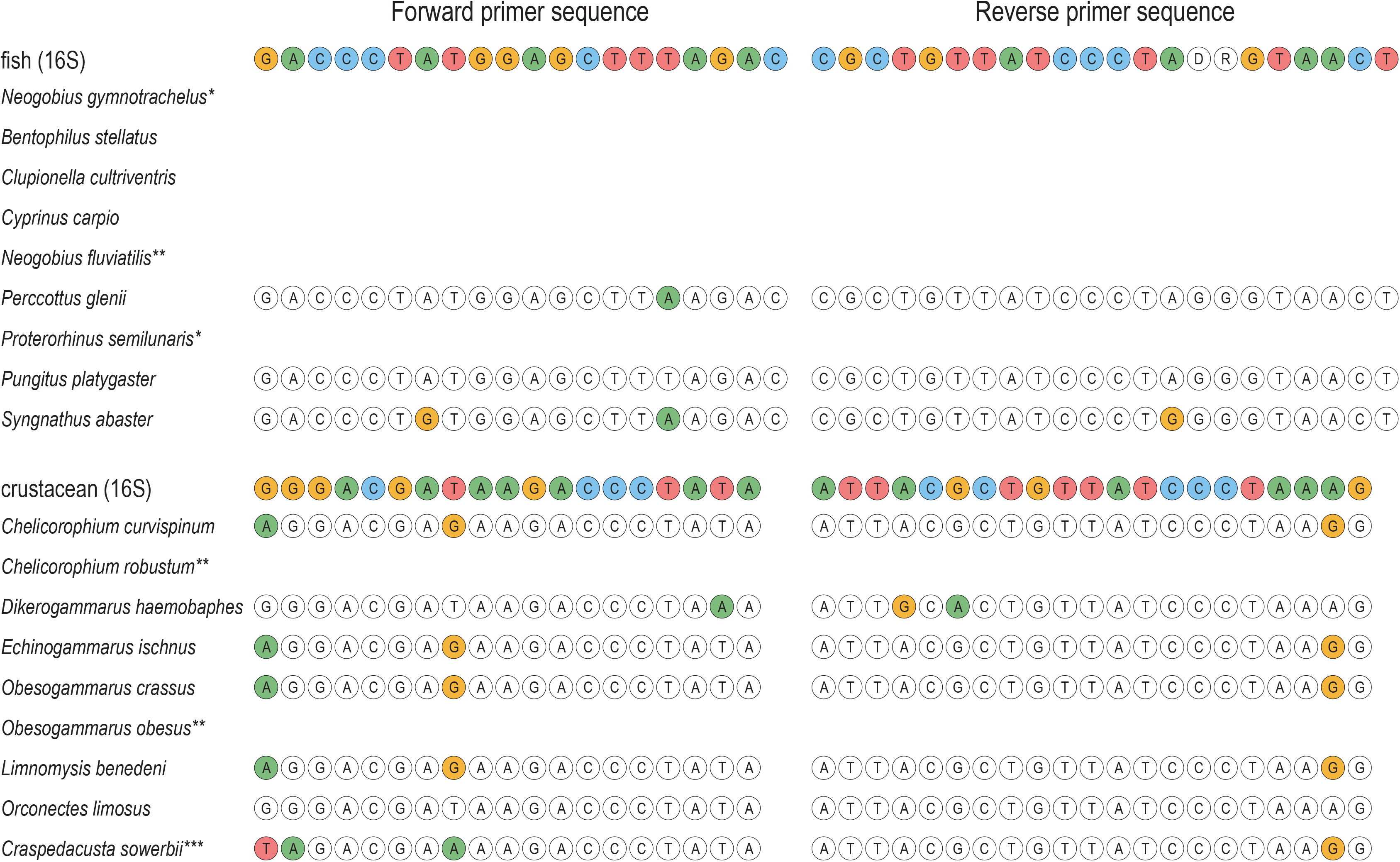
In silico PCR analysis identifying the completeness of the reference database and mismatches in the forward and reverse primer binding site for the fish (16S) and crustacean (16S) assay. Mismatches in the primer binding sites are indicated by colored circles. * denotes the presence of a reference sequence without primer-binding regions, ** denotes newly barcoded species, and *** denotes species only detected by the eDNA survey.

Two of the nine NIS picked up in our *in silico* PCR did not display mismatches in the primer-binding sites, including one fish (*Pungitus platygaster*) and one invertebrate (*Faxonius limosus*). One fish (*Perccottus glenii*) displayed a single mismatch in the primer-binding sites, while the remaining NIS displayed three mismatches. Mismatches in the forward primer were found in the 5’ end, while mismatches in the reverse primer were found in the 3’ end for crustacean NIS, potentially influencing amplification efficiency for this taxonomic group.

### Locating the range of invasion – fish NIS

Our ichthyological survey detected seven of the nine NIS on the Dnieper River (site NZ) near the southern border with Ukraine, the entry point of invasion (FIGURE 3). Four NIS (*Benthophilus gymnotrachelus*, *Clupionella cultriventris*, *Pungitius platygaster*, and *Syngnathus abaster*) were only found at site NZ in low abundance. The most widely distributed NIS according to our ichthyological survey was *Neogobius fluviatilis*, followed by *Neogobius gymnotrachelus*, and *Proterorhinus semilunaris*. All three species were detected at multiple sites throughout the southern region of Belarus on the Dnieper and Pripyat rivers. Two NIS were not detected at site NZ. *Perccottus glenii* was only detected at site PN, while *Cyprinus carpio* was detected in the two most northern sites, i.e., site ZD and site DVD. Non-indigenous fish were found at eleven sites, excluding one site on the Neman River (site N). The highest number of NIS fish (seven species) was detected on the Dnieper River (site NZ), followed by 5 NIS fish on the Pina River (site PP). Highest abundance of NIS fish (13.59%) was detected on the Dnieper River (site NZ), followed by 11.74% and 11.68% on the Pripyat River (site PN) and Dnieper-Bug canal (site DBD), respectively.

**Figure 3:**
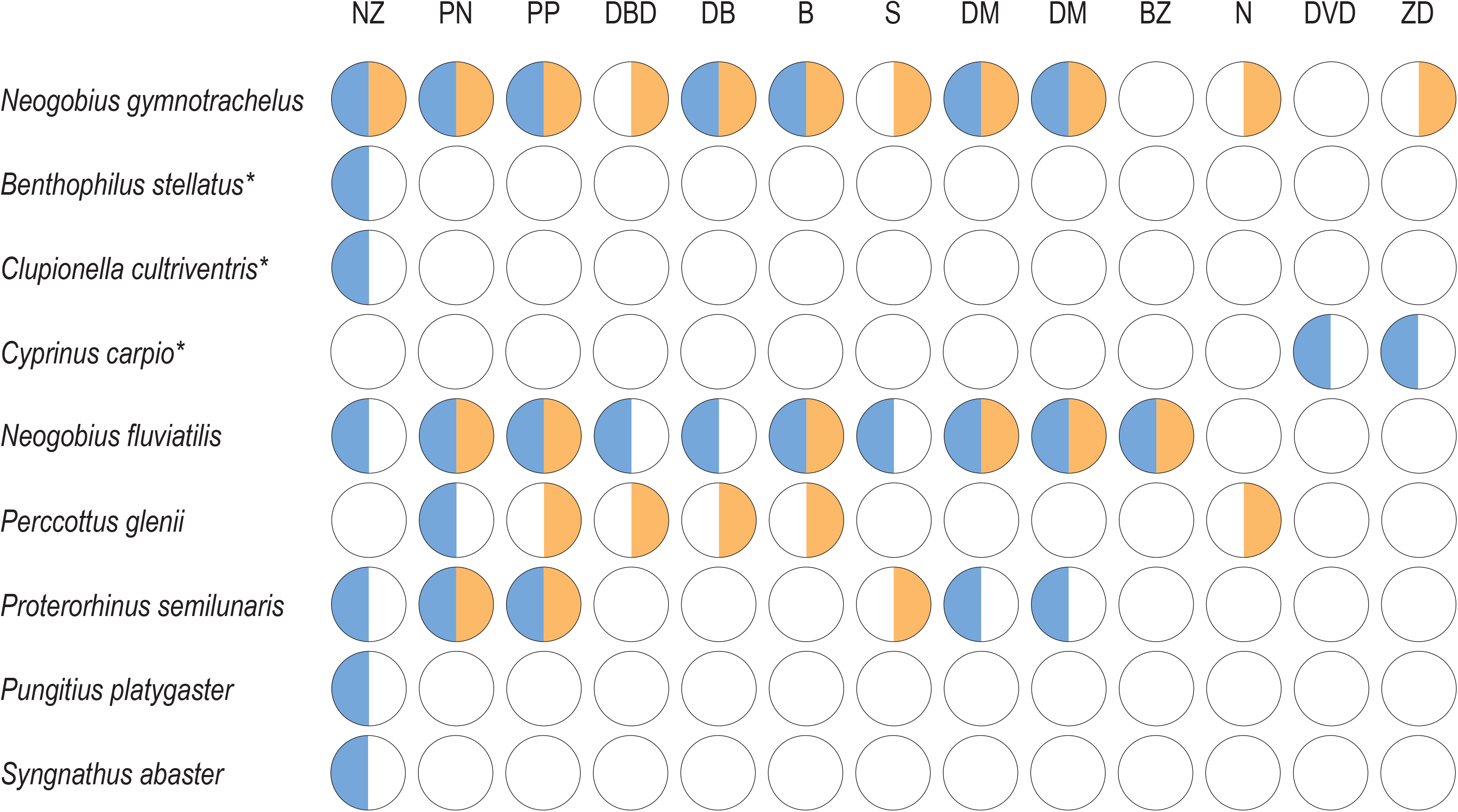
Site-specific non-indigenous fish detection for the ichthyological and eDNA survey. Positive detection is indicated by colored circles, with a positive detection for the eDNA survey in orange and positive detection for the ichthyological survey in blue. * denotes NIS without a reference barcode sequence. Sampling site notation follows the abbreviations of Table 1.

Our fish eDNA metabarcoding survey detected four of the six non-indigenous fish species with a reference barcode, while failing to detect two fish NIS, i.e., *Pungitius platygaster* and *Syngnathus abaster*, both detected in low abundance at a single site by our ichthyological survey (FIGURE 3). The most widely distributed NIS according to our eDNA survey was *Neogobius gymnotrachelus*, followed by *Neogobius fluviatilis*, *Perccottus glenii*, and *Proterorhinus semilunaris*. All NIS were detected in the southern region of Belarus and were detected further north compared to our ichthyological survey (FIGURE 3). Non-indigenous fish were found at eleven sites, excluding one site on the Zapadnaya Dvina River (site DVD). All non-invasive fish were detected on the Pina River (site PP), while three NIS were detected on the Muchavets River (site B) and Pripyat River (site PN). Based on relative number of reads, highest abundance of NIS fish (40.25%) was detected on the Dnieper River (site NZ), followed by 27.54%, 21.85%, and 20.56% on the Dnieper River (site DM), Pina River (site PP), and Dnieper-Bug canal (site DBD), respectively.

### Locating the range of invasion – invertebrate NIS

Non-invasive macro-invertebrates were recorded at all twelve sites. The highest number of NIS (seven species) was detected on the Pripyat River (site PN), followed by six NIS on the Dnieper River (site NZ) and five NIS on the Muchavets River (site B). Highest abundance of NIS (405 individuals) was detected on the Pripyat River (site PN), followed by 81 and 80 individuals on the Dnieper River (site NZ) and Sozh River (site S), respectively (SUPPLEMENT 3). According to the hydrobiological survey, the gravel snail (*Lithoglyphus naticoides*) is most widely distributed with a positive detection at ten sites, followed by *Dikerogammarus haemobaphes* with a positive detection at seven sites (FIGURE 4). The highest abundant NIS were *Obesogammarus crassus* and *Lithoglyphus naticoides*, with 331 and 174 detections, respectively. The least widely distributed NIS with a detection at a single site (site PN) was *Obesogammarus obesus*, followed by *Faxonius limosus*, *Limnomysis benedeni*, *Echinogammarus ischnus*, and *Chelicorophium robustum*, which were detected at two sites. The least abundant NIS were *Faxonius limosus*, *Obesogammarus obesus*, and *Chelicorophium robustum*, with two, two, and three detections, respectively (SUPPLEMENT 3).

**Figure 4:**
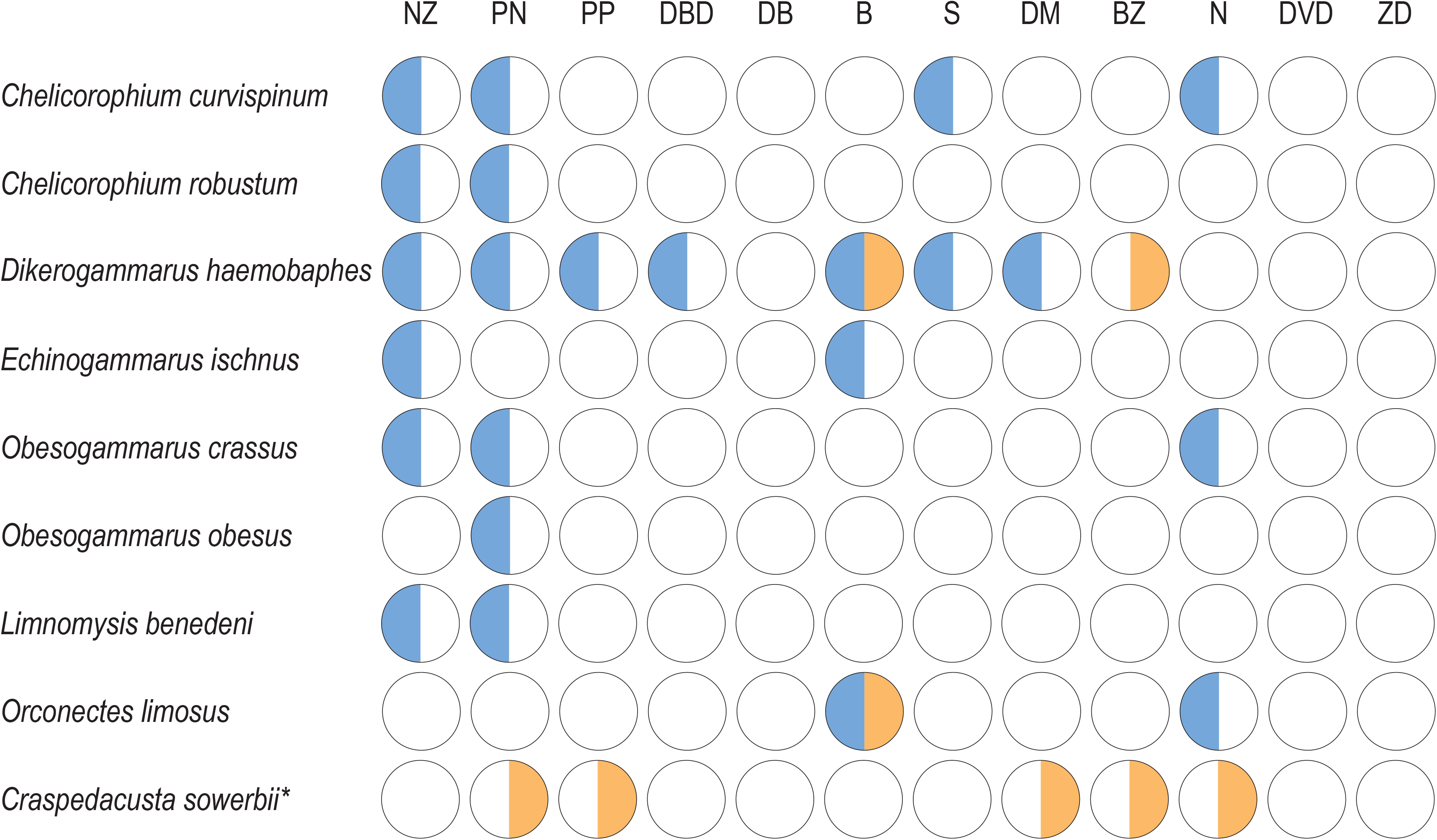
Site-specific non-indigenous macro-invertebrate detection for the hydrobiological and eDNA survey. Positive detection is indicated by colored circles, with a positive detection for the eDNA survey in orange and positive detection for the hydrobiological survey in blue. * denotes NIS detected solely by the eDNA survey. Sampling site notation follows the abbreviations of Table 1.

Our eDNA metabarcoding survey detected only two of the eight non-indigenous macro-invertebrates with a reference barcode, i.e., *Dikerogammarus haemobaphes* and *Faxonius limosus* (FIGURE 4). For both invasive species, the eDNA survey obtained a positive detection in a reduced number of sites compared to the hydrobiological survey. However, the eDNA survey detected *Dikerogammarus haemobaphes* at one additional site on the Berezina river (site BZ). The eDNA survey detected an additional NIS, a freshwater jellyfish (*Craspedacusta sowerbii*), not detected by the hydrobiological survey. *Craspedacusta sowerbii* is a known European invader, previously not yet reported in Belarussian rivers and lakes, but was found in several artificial water bodies in the Pripyat and Muchavets River basins by local people. According to the eDNA survey, the freshwater jellyfish (*C. sowerbii*) was most widely distributed with a positive eDNA signal at five sites, while *Dikerogammarus haemobaphes* was only detected in two out of seven sites compared to the hydrobiological survey and the spinycheek crayfish (*F. limosus*) was only detected at a single site on the Muchavets River (site B) compared to two sites in the hydrobiological survey (FIGURE 4).

## Discussion

In this study, we evaluated eDNA metabarcoding as an alternative survey method for the simultaneous detection of non-indigenous species in riverine systems. Our results provide compelling evidence that eDNA metabarcoding on DNA extracted from surface water samples can be implemented for aquatic NIS monitoring in freshwater environments to explore the range of invasion (Holman et al. 2019; van den Heuvel-Greve et al. 2021). The complexity of the DNA signal from environmental samples, furthermore, enables the detection of unexpected NIS (e.g., the freshwater jellyfish *Craspedacusta sowerbii*) that would not be targeted in established traditional monitoring methods, such as the ichthyological and hydrobiological surveys currently conducted in Belarus.

By extending the range of invasion for two non-indigenous fish species, we document the potential of eDNA metabarcoding from low-volume surface-water samples to detect aquatic NIS at an early stage of invasion. Incorporating eDNA metabarcoding surveys into established conservation programs will, thereby, increase the chance for successful eradication of aquatic NIS (Reaser et al. 2020). However, due to the lack of physical evidence, i.e., no specimens sighted at these sampling sites, increased monitoring is required in the future to validate the eDNA metabarcoding results. Once validated, eDNA metabarcoding surveys could be used as a guide for increased monitoring effort at specific locations.

The invasion range extension noted by eDNA metabarcoding might be attributed by increased sensitivity over the ichthyological survey, as reported in several comparative studies between traditional and active eDNA surveys (Ardura et al. 2015; Dougherty et al. 2016; Simpfendorfer et al. 2016). However, the multiple recorded false-negative detections for the eDNA survey in this study questions an overall increased sensitivity for aquatic organisms (FIGURE 3). Furthermore, while reporting the limit of detection (LOD) and quantification (LOQ) has been suggested for active eDNA surveys (Klymus et al. 2020), sensitivity estimates for passive eDNA surveys are difficult to assess, as amplification efficiency will be highly dependent on the residing community (Krehenwinkel et al. 2017; Kelly et al. 2019). Alternatively, DNA transport following river flow from the source population or individual could potentially lead to false-positive detection (Roussel et al. 2015). However, the spatial resolution of eDNA has previously been reported in the range of meters to kilometers (Deiner and Altermatt 2014; Deiner et al. 2016; Rice et al. 2018), while our sampling sites are situated tens/hundreds of kilometers apart (FIGURE 1). DNA could be further transported, however, by fecal contamination from predators such as migrating birds (Merkes et al. 2014). To limit issues surrounding false-positive species detection through DNA transport, environmental RNA (eRNA) metabarcoding might be a suitable alternative due to the less stable nature of the molecule (Pochon et al. 2017; Zaiko et al. 2018; von Ammon et al. 2019).

One additional aquatic NIS was detected by the crustacean (16S) assay, i.e., the freshwater jellyfish *Craspedacusta sowerbii*. While both traditional monitoring methods employed in Belarus are field standards, they target specific taxonomic groups that do not cover invertebrate organisms residing in the water column. Environmental DNA metabarcoding, on the other hand, takes advantage of the complexity of the DNA signal from environmental samples, facilitating the detection of unexpected NIS provided a reference barcode is available. *Craspedacusta sowerbii* natively inhabits freshwater bodies of Eastern Asia (Jankowski et al. 2008) and was first recorded in Europe (United Kingdom) in 1880 (Boothroyd et al. 2002) and 1901 in mainland Europe (Lytle 1960). While *Craspedacusta sowerbii* has been recorded in neighboring countries, such as Ukraine and Poland (Arbačiauskas and Lesutienė 2005; Didžiulis and Zurek 2013), this is the first record of the freshwater invasive jellyfish in Belarus. The role of freshwater jellyfish in food webs, as well as their impact on local aquatic communities still remains insufficiently studied (Dumont 1994). While the direct impact for Belarussian riverine communities might be restricted to the predation of fish eggs (Dumont 1994), *Craspedacusta sowerbii* might secondarily enhance the spread of the non-indigenous spinycheek crayfish (*Faxonius limnosus*). The spinycheek crayfish actively predates on this freshwater jellyfish under laboratory conditions (Dodson and Cooper 1983). The presence of *Craspedacusta sowerbii* might, therefore, increase the available food source of this alien crayfish. Future surveys and monitoring are required to obtain a specimen for confirmation of this NIS in Belarussian waters and to determine the impact of *Craspedacusta sowerbii* on the native riverine communities.

Our ichthyological survey picked up an increased number of non-indigenous fish species compared to the eDNA metabarcoding survey. Three NIS missed by our eDNA survey do currently not have a reference barcode available on public databases for the 16S rRNA gene targeted by the fish (16S) assay (FIGURE 2). With eDNA metabarcoding relying on species-identification through reference barcodes, one major limitation is the incompleteness of the reference database (Hestetun et al. 2020). Although cytochrome c oxidase subunit I (COI) boasts the most complete reference database for eukaryotes to date (Collins et al. 2019), primer assay efficiency issues and an increased risk of false-negative detections have been reported for COI in eDNA metabarcoding studies (Collins et al. 2019; Hajibabaei et al. 2019; Jeunen, Knapp, et al. 2019). Therefore, primer assay choice will, currently, be a trade-off between completeness of the reference database and assay amplification efficiency and specificity (Krehenwinkel et al. 2017; Kelly et al. 2019). Ultimately, continuous barcoding of multiple genetic markers or complete mitogenomes will be essential to exploit the true potential of eDNA metabarcoding in the future (Collins et al. 2019).

Two non-indigenous fish species, detected at low abundance in the southernmost site (site NZ) by the ichthyological survey, failed to be detected by the eDNA survey, i.e., the black-striped pipefish (*Syngnathus abaster*) and the southern nine spine stickleback (*Pungitius platygaster*). These false-negative detections could indicate a need for increased sampling effort and processing volume (Cantera et al. 2019). While Sterivex filtration is frequently used in eDNA studies, the limit in volume passing through the filter until clogging might enhance an increased risk of false-negative detections (Li et al. 2018). Since eDNA degrades during long transport times to dedicated PCR-free laboratory facilities and on-site vacuum filtration might not be feasible due to a lack of a power source and clean workspace, passive filtration (Bessey et al. 2021) and filter-feeding organisms (Mariani et al. 2019) could provide suitable alternatives to increase the volume processed per sample and limit false-negative detections. Another explanation for the false-negative detections might be related to the high spatial and temporal resolution of eDNA (Beentjes et al. 2019; Brys et al. 2020). Since both monitoring methods were not conducted simultaneously, eDNA samples might have been collected when both species were not present at the time of sampling, indicating increased sampling effort is needed for the reliable detection of migrating or non-established species. Both false-negative detections for low-abundant fish species might also be attributed to DNA degradation during the long transportation time of samples from Belarus to New Zealand, extending the time between sample collection and DNA extraction. While filter storage in Longmire’s buffer has been shown to effectively preserve eDNA over short periods of time (Renshaw et al. 2015), immediate DNA extraction is preferred (Kumar et al. 2020).

Environmental DNA obtained from surface water failed to reliably detect non-indigenous, sediment-living macro-invertebrates. Previous studies reported different eDNA signals obtained from various substrates originating from the residing community (Turner et al. 2015; Koziol et al. 2019). Furthermore, different eDNA signals have been obtained from different depths in the water column in stratified conditions (Jeunen, Lamare, et al. 2019; Littlefair et al. 2020). Additionally, the macro-invertebrate eDNA signal retrieved from the water column in riverine systems has been shown to differ from the detected diversity from benthic bulk samples, indicating aqueous eDNA might not be effective at detecting benthic taxa (Gleason et al. 2020). The results from the crustacean (16S) assay in this study corroborate these findings, as the majority of detected taxa consisted of aquatic and aquatic-associated invertebrates, such as copepods and dragonflies (SUPPLEMENT 5). Although eDNA metabarcoding has the potential to aid monitoring efforts in the early detection of NIS, data obtained from a single substrate could be insufficient and increase the risk of false-negative species detection. A more substantial sampling strategy incorporating multiple substrates is, therefore, recommended.

An increase in detection probability for non-indigenous species would further enhance the power of eDNA metabarcoding as a survey method for the early detection of invasive taxa. For this initial study, we made use of two previously designed and published primer assays. Although our eDNA survey successfully detected a myriad of NIS and extended the invasion range further north for two aquatic NIS, the *in silico* PCR analysis identified multiple mismatches in the forward and reverse primer-binding sites for the majority of target NIS (FIGURE 2), potentially influencing the amplification efficiency in a negative manner (Stadhouders et al. 2010). Future assay optimization could increase amplification efficiency and, hence, the detection probability for target non-indigenous species. Furthermore, inclusion of blocking primers to exclude DNA signals originating from the host organism in dietary studies (Robeson II et al. 2018) or highly abundant species in environmental monitoring has shown to increase the detection probability for rare species and reduce the minimum required sampling effort (Wilcox et al. 2014; Rojahn et al. 2021).

## Conclusions

With this comparative experiment, we provide evidence for the potential of eDNA metabarcoding to record the invasion range of multiple non-indigenous species in an accurate, cost-effective, and time-efficient manner. However, current drawbacks, including incomplete reference databases (Hestetun et al. 2020), reliance on amplification methods for species detection (Wilcox et al. 2018), detection probability (Rojahn et al. 2021), and sampling effort (Li et al. 2018) are limiting the true potential of this revolutionary monitoring method. Nevertheless, eDNA metabarcoding has the potential to aid monitoring efforts in the early detection of NIS and guide future monitoring efforts to specific locations. Furthermore, by taking advantage of the complex DNA signal contained within environmental samples, eDNA metabarcoding increases the chance to detect unexpected NIS. We, therefore, recommend the implementation of eDNA metabarcoding surveys alongside traditional approaches to increase the probability of early NIS detection and, hence, facilitate successful eradication efforts and minimize ecological impacts.

## Acknowledgements

We would like to thank Sara Ferreira and Joanne Gillum for their assistance with laboratory work. This study has been partly supported by the Belarusian Republican Foundation for Fundamental Research (Grant № B19MS-026) – TL.

