## Supplement 1 and 4 for "Environmental DNA (eDNA) metabarcoding surveys extend the range of invasion for non-indigenous freshwater species in Eastern Europe"

Supplement 1: Metabarcoding qPCR assays

and the respective primer sets used for biodiversity detection.

| Metabarcoding assay | Primer set | Target | Primer sequence | Gene | Amplicon length (bp) | Reference | Assay T_m_ (°C) |
| --- | --- | --- | --- | --- | --- | --- | --- |
| fish (16S) | Fish16SF/D | Fish | 5’-GACCCTATGGAGCTTTAGAC-3’ | 16S rRNA | ~200 | (Berry et al., 2017) | 54 |
|  | 16S2R |  | 5’-CGCTGTTATCCCTADRGTAATC-3’ |  |  |  | 51 |
| crustacean (16S) | Crust16S_F(short) | Crustacea | 5’-GGGACGATAAGACCCTATA-3’ | 16S rRNA | ~170 | (Berry et al., 2017) | 51 |
|  | Crust16S_R(short) |  | 5’-ATTACGCTGTTATCCCTAAAG-3’ |  |  |  |  |

*Supplement 2: Relative fish abundances as observed by the ichthyological survey. Scientific and common names are given in the respective columns. Non-indigenous species are identified by “Yes” in the “Invasive” column. Values indicate the relative abundance at a given sampling site, while “+” indicates a positive detection at sampling sites “ZD” and “DVD”. Sampling site notation follows the abbreviations of Table 1. Data can be found in the “supplement_2_ichthyological_survey.csv” file.*

*Supplement 3: Macro-invertebrate abundances as observed by the hydrobiological survey. Scientific names are given in the respective column. Values indicate the number of individuals at a given sampling site. Sampling site notation follows the abbreviations of Table 1. Data can be found in the “supplement_3_hydrobiological_survey.csv” file.*

*Supplement 4: Rarefaction curves for each metabarcoding assay (fish (16S); crustacean (16S)) per habitat for each sampling site. Number of taxa are indicated on the y-axis and number of reads on the x-axis. Sampling site notation follows the abbreviations of Table 1.*

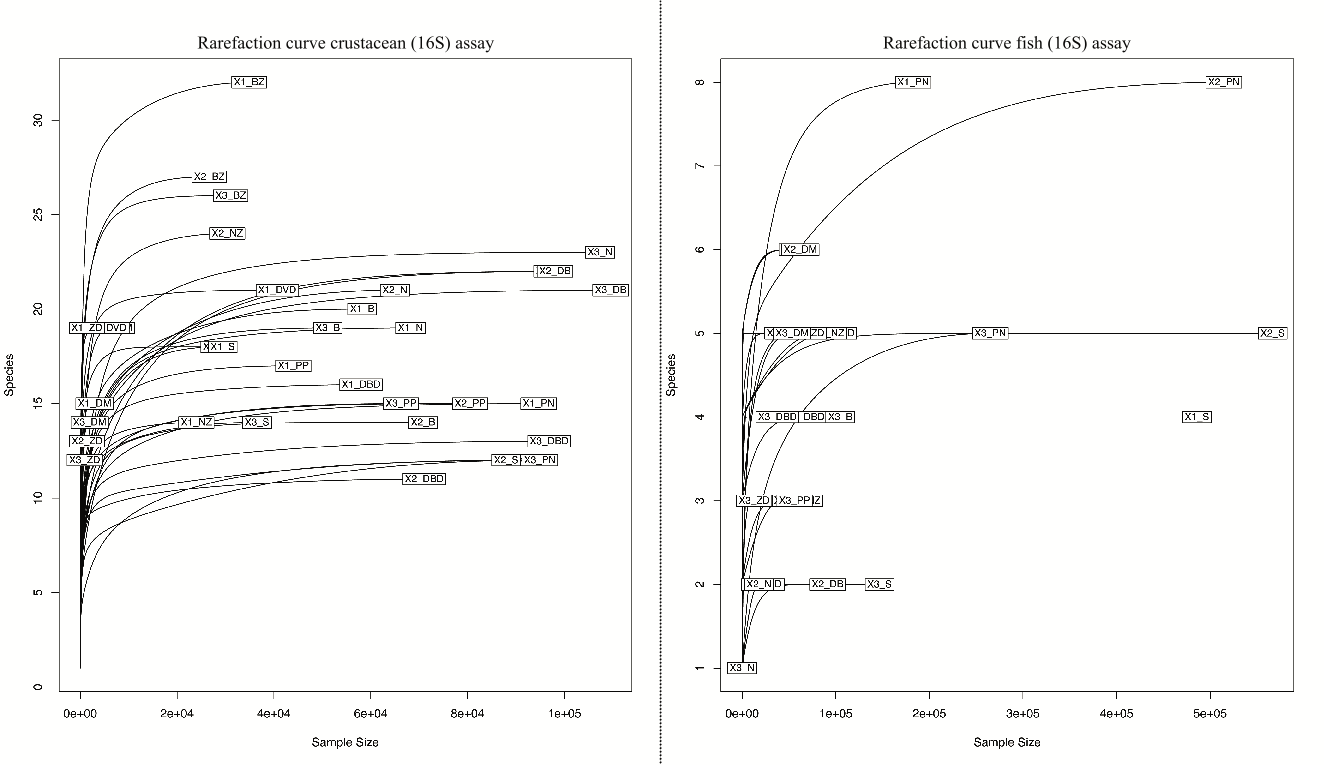

*Supplement 5: Environmental DNA detections from the passive surveillance for both metabarcoding assays, i.e., fish (16S) and crustacean (16S). Scientific and common names are given in the respective columns. Values indicate the number of reads assigned to each taxonomic unit for a given sampling site. Sampling site notation follows the abbreviations of Table 1. Data can be found in the “supplement_5_eDNA_survey.csv” file.*
